# High-Speed Imaging Reveals the Bimodal Nature of Dense Core Vesicle Exocytosis

**DOI:** 10.1101/2021.11.01.466835

**Authors:** Pengcheng Zhang, David Rumschitzki, Robert H. Edwards

## Abstract

During exocytosis, the fusion of secretory vesicle with plasma membrane forms a pore that regulates release of neurotransmitter and peptide. Heterogeneity of fusion pore behavior has been attributed to stochastic variation in a common exocytic mechanism, implying a lack of biological control. Using a fluorescent false neurotransmitter (FFN), we imaged dense core vesicle (DCV) exocytosis in primary mouse adrenal chromaffin cells at millisecond resolution and observed strikingly divergent modes of release, with fast events lasting <30 ms and slow events persisting for seconds. Dual imaging of slow events shows a delay in the entry of external dye relative to FFN release, suggesting exclusion by an extremely narrow pore <1 nm in diameter. Unbiased comprehensive analysis shows that the observed variation cannot be explained by stochasticity alone, but rather involve distinct mechanisms, revealing the bimodal nature of DCV exocytosis. Further, loss of calcium sensor synaptotagmin 7 increases the proportion of slow events without changing the intrinsic properties of either class, indicating the potential for independent regulation. The identification of two distinct mechanisms for release capable of independent regulation suggests a biological basis for the diversity of fusion pore behavior.

## INTRODUCTION

Regulated exocytosis provides a mechanism to control the timing of signals important for development, physiology and behavior. The latency from stimulus to membrane fusion, or synchrony, makes an important contribution to the time course of signaling. In neurons, synaptic vesicle exocytosis within milliseconds of stimulation confers the synchronous release of neurotransmitter essential for brain function (Kaeser and Regehr, 2014). In endocrine cells, release from dense core vesicles (DCVs) exhibits a biphasic time course with initial, rapid release followed by slow, sustained secretion (Parsons et al., 1995; Voets et al., 1999). The biphasic release of insulin from pancreatic β-cells plays a critical role in glucose homeostasis, and loss of the first phase is considered a hallmark of type 2 diabetes (Rorsman and Renstrom, 2003). The initial rapid phase of release involves a readily releasable pool of vesicles at the plasma membrane, whereas the slower phase requires activation of additional vesicles through docking or priming. The variation in synchrony within individual cells is thus thought to reflect vesicles at different steps along a common pathway to exocytosis.

After membrane fusion, release from individual vesicles also varies in time course. Upon fusion, a channel known as the fusion pore connects the vesicle lumen to the extracellular space (Breckenridge and Almers, 1987). The properties and behavior of the fusion pore affect the time course of release in multiple ways. Pore size influences the rate of release: narrow pores slow secretion and wide pores accelerate (Bao et al., 2018; Chow et al., 1992; Shin et al., 2018). The fusion pore can also act as a molecular sieve that differentially affects the release of cargo based on their size or charge (Fulop et al., 2005; Perrais et al., 2004). In addition, the fusion pore can dilate, leading to the discharge of all contents and vesicle collapse into the plasma membrane. Alternatively, the fusion pore can reseal, resulting in a form of exocytosis known as ‘kiss-and-run’ that preserves the vesicle for additional rounds of release (Alabi and Tsien, 2013), thus enabling repeated response to a prolonged signal. The fusion pore influences release from DCVs in many cell types including neurons as well as endocrine cells (Breckenridge and Almers, 1987; Logan et al., 2017; Xia et al., 2009). The fusion pore also exhibits remarkable heterogeneity within individual cells. Since a single pathway to exocytosis was inferred from the analysis of synchrony, the heterogeneity in pore behavior is attributed to stochastic variation, which implies a lack of biological control.

Previous work has characterized the diversity of fusion pore behavior by classifying exocytic events based on specific characteristics. For example, DCV exocytosis has been imaged using fluorescent reporters, and fluorescence time trace classified based on the shape of the curve (Chiang et al., 2014; Logan et al., 2017). Similarly, the amperometric measurement of catecholamine release shows spikes indicating rapid pore expansion and ‘feet’ suggestive of narrow pores, and these are analyzed separately (Zhou et al., 1996). The properties of each class are thus quantified using different metrics, making it difficult to determine whether the classes differ in mechanism or reflect only stochastic variation.

In addition, the available methods reveal only specific features of the exocytic process. Amperometry has high temporal resolution but lacks spatial information and does not monitor DCV behavior before fusion. Diffusion also makes the measurement of event kinetics sensitive to the distance between release site and electrode (Albillos et al., 1997). In contrast, imaging by total internal reflection fluorescence (TIRF) microscopy provides high spatial resolution and can monitor DCV behavior before exocytosis. However, the weak emission of conventional fluorophores requires long exposure times, limiting temporal resolution. The inability to simultaneously monitor multiple features of an individual exocytic event produces a fragmentary description, potentially failing to capture a relationship between features that has important implications for mechanism.

We now describe a method for imaging that detects secretory vesicles before fusion and monitors release at high temporal resolution, providing a holistic picture of exocytosis that enables us to characterize the diversity of fusion pore behavior. We identify two distinct modes of fusion pore behavior, showing that stochasticity alone cannot account for the diversity.

## RESULTS

### Imaging Exocytic Events at Millisecond Resolution

Fluorescent false neurotransmitters (FFNs) are fluorescent catecholamine analogues recognized as substrates by the vesicular monoamine transporter (VMAT) and hence concentrated into neurosecretory vesicles (Gubernator et al., 2009; Hu et al., 2013). The pH-insensitive FFN206 (Figure 1—figure supplement 1A,B), which is not quenched by the acidic lumenal environment (Figure 1—figure supplement 1C,D), enables us to detect vesicles before fusion. When loaded into primary adrenal chromaffin cells, FFN206 accumulates in chromaffin granules, presumably due to the catalytic activity of VMAT, conferring a high signal-to-noise ratio that enables imaging at millisecond resolution. Using TIRF microscopy, we observe a punctate distribution of FFN-loaded vesicles illuminated by the evanescent field (Figure 1A), presumably vesicles docked at the plasma membrane. Stimulation produces a large number of transient events that we imaged at an acquisition rate of 800 Hz (Figure 1B–D, Figure1—figure supplement 2, and Figure 1—video 1). For each event, we observe an increase in fluorescence suggestive of dye approaching the TIRF plane at the coverslip surface due to exocytosis. To focus on events originating within the TIRF plane, we excluded those with diffuse fluorescence appearing simultaneously over a large area, and counted only events that begin as a point source. Consistent with an exocytic mechanism, all FFN events require stimulation (Figure 1—video 1) and external Ca^++^ (Figure 2A). Cleavage of v-SNAREs by tetanus toxin (McMahon et al., 1993; Schiavo et al., 1992) abolishes the FFN response (Figure 2B), further supporting an exocytic origin for all events.

**Figure 1.**
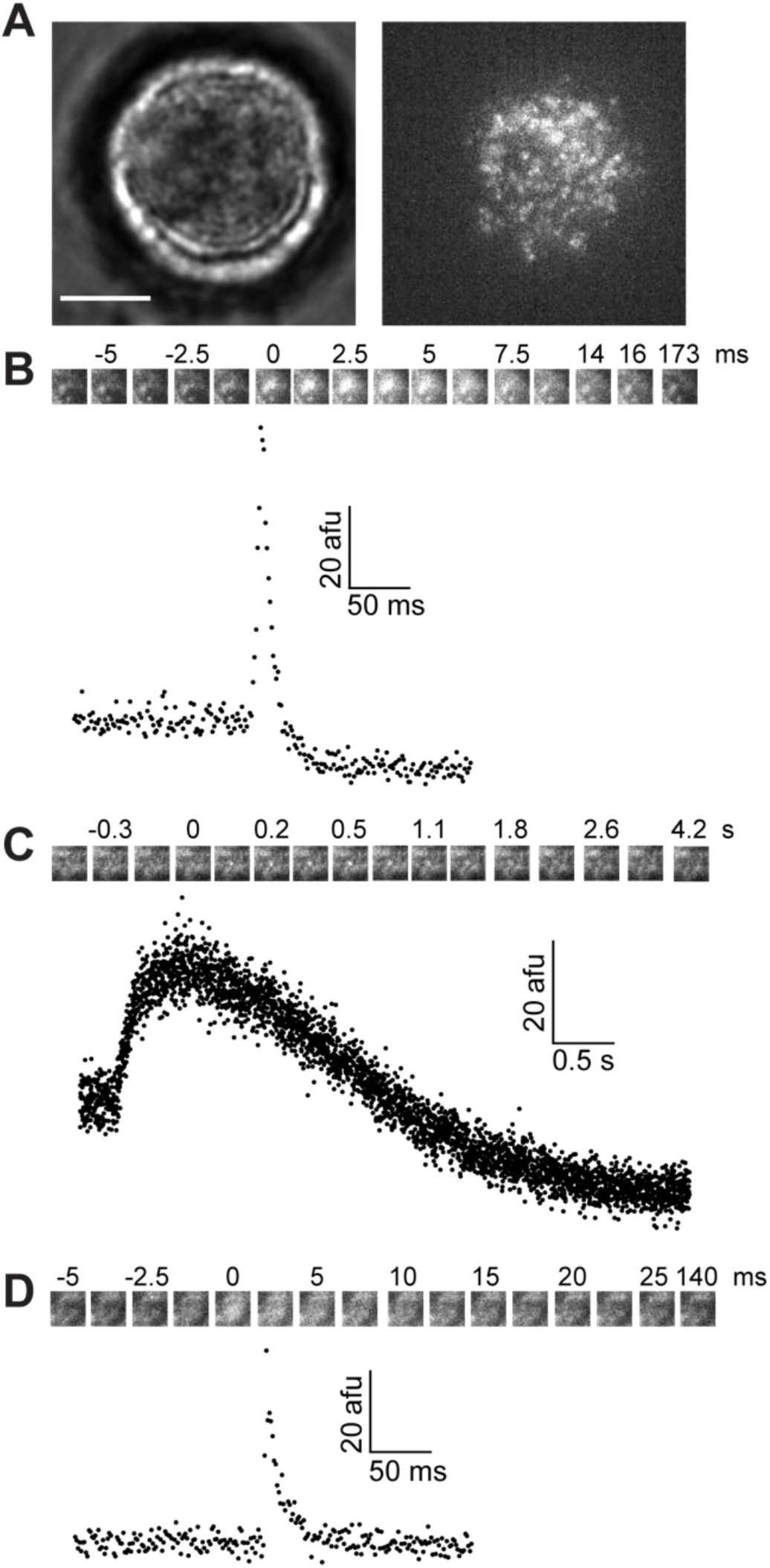
High frequency imaging reveals distinct classes of exocytic event. (**A**) Micrographs of an isolated primary mouse adrenal chromaffin cell, illuminated by bright field (left) or total internal reflection, revealing FFN puncta (right). Scale bar: 5 µm. (**B–D**) Fluorescence time traces showing fast (**B**) and slow (**C**) release of FFNs with reduced post-event baseline (indicating docked vesicles), as well as rapid release with unchanged baseline (no detectable docked vesicle) (**D**). Insets in each panel show select frames of the corresponding event, marked by the time relative to initial FFN signal increase (time 0). **Figure 1—figure supplement 1.** FFN properties. **Figure 1—figure supplement 2.** Sample fluorescence time traces of FFN release events. **Figure 1—figure supplement 3.** Tightness of cell attachment does not affect FFN release kinetics. **Figure 1—video 1.** High-speed imaging of FFN release from an isolated primary mouse adrenal chromaffin cell in culture. **Figure 1—video 2.** Fast FFN release from a docked vesicle. **Figure 1—video 3.** Fast FFN release without prior detectable docked vesicle. **Figure 1—video 4.** Slow FFN release from a docked vesicle. **Figure 1—video 5.** Dual imaging of FFN release and BDNF-pHluorin exocytosis.

**Figure 2.**
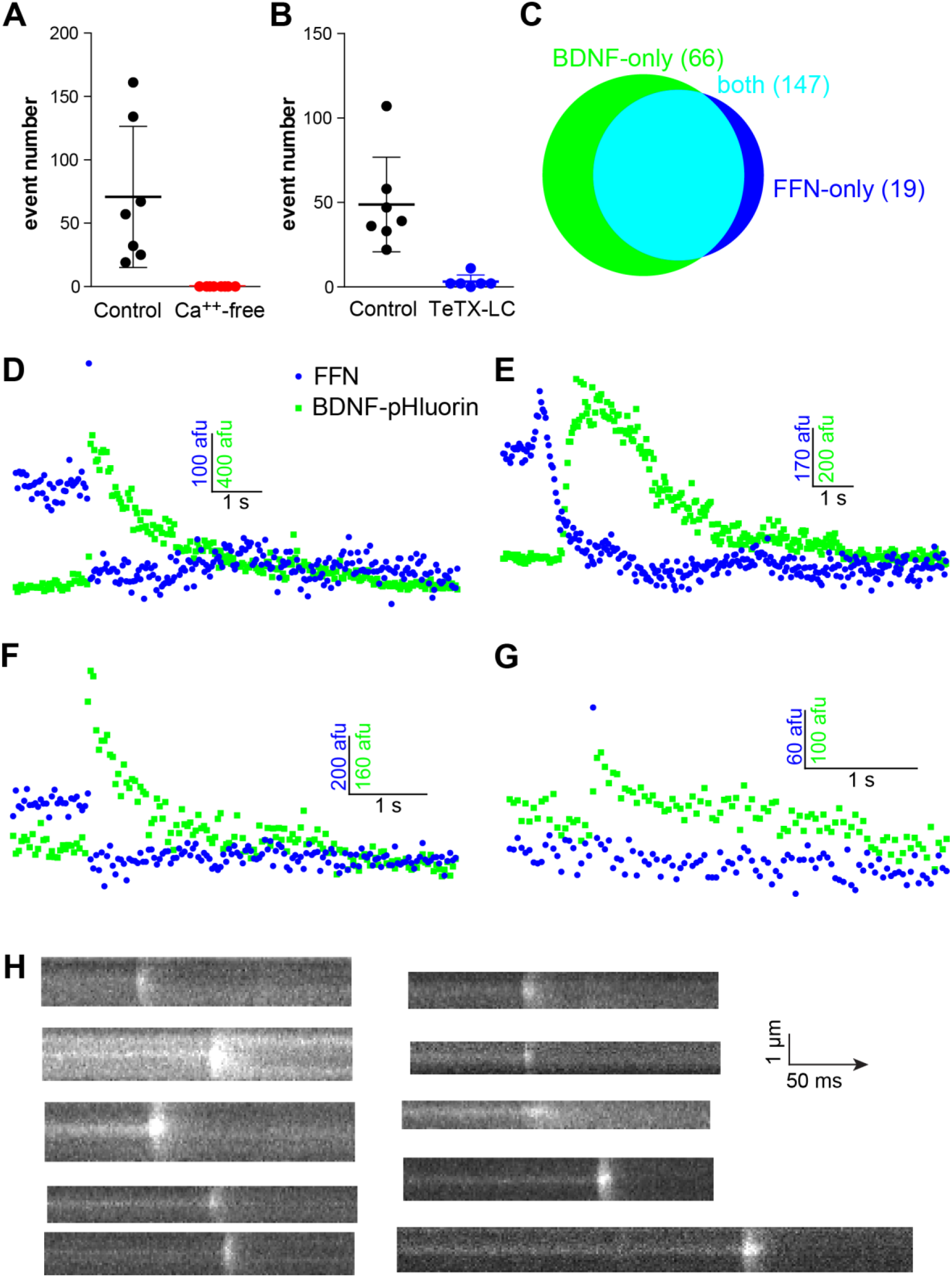
FFN release is exocytic in nature. (**A**) Removal of Ca^++^ from the external solution abolishes FFN release from chromaffin cells. (**B**) Lentivral transduction of tetanus toxin light chain (TeTX-LC) abolishes FFN release. Error bars indicate standard deviation. (**C**) Quantitative Venn diagram shows the vast majority of FFN release events coincide with unquenching of BDNF-pHluorin. (**D–G**) Fluorescence time traces show that unquenching of BDNF-pHluorin (green) follows FFN release (blue) for both fast (**D** and **F**) and slow events (**E**) with reduced post-event baseline, as well as for release with an unchanged post-event baseline (**G**). The prolonged exposure time required for dual imaging of BDNF-pHluorin and FFN limited capture of the FFN event to a single (**D** and **G**) or no (**F**) frame. (**H**) Kymographs showing the spread of FFN following rapid release.

The majority of FFN events involve a characteristic, brief spike in fluorescence (Figure 1B, Figure 1—figure supplement 2A, and Figure 1—video 2) followed by dispersion (Figure 2H). This is consistent with bulk release of vesicular FFN into the extracellular space (where the evanescent field is exponentially stronger) followed by rapid diffusion away from the release site. These fast events presumably reflect release through a large fusion pore. Almost all events result in reduction of baseline fluorescence (Figure 1B,C, and Figure 1—figure supplement 2A,B), which we interpret as the release of all FFN from vesicles docked at the plasma membrane. However, a small fraction (1–10%) of exocytic events do not involve a change in baseline fluorescence (Figure 1D, Figure 1—figure supplement 2C, and Figure 1—video 3). These might reflect release from vesicles that arrived from the cell interior and fused with essentially no delay. Alternatively, they may reflect release from docked vesicles lying just above the TIRF plane. We thus assessed the distance between the cover glass surface and plasma membrane, using the fluorescence intensity of external cell-impermeant Alexa dye in the evanescent field as a proxy for the tightness of attachment (Figure 1—figure supplement 3A–C). We found no difference in attachment tightness at the sites of release regardless of whether docked vesicles were detected or not (Figure 1— figure supplement 3D). The peak amplitude of fast events without an observed docked vesicle was slightly higher than that of events with docked vesicles (Figure 1—figure supplement 3E), arguing against the possibility that events without an observed docked vesicle are further from the cover glass. However, significant variation in Alexa fluorescence was also observed in cell-free regions (Figure 1—figure supplement 3B), indicating that the unevenness of the cover glass surface is non-negligible, thereby complicating our estimates of tightness of attachment. Thus, we cannot exclude the possibility of fusion by a docked vesicle just outside the TIRF plane, which is consistent with previous work showing that synaptic vesicles fuse ∼60 ms after docking (Chen et al., 2013), and suggesting that DCVs may require more time (Steyer et al., 1997).

A substantial fraction of events persists from hundreds of milliseconds to seconds, exhibiting highly variable fluorescence traces without a recognizable spike (Figure 1C, Figure 1— figure supplement 2B, and Figure 1—video 4). These slow events presumably reflect the efflux of FFN through a narrow fusion pore that limits diffusion of even a small molecule (Shin et al., 2018). Amperometric recording of catecholamine release has identified ‘stand-alone feet’ that reflect the slow efflux of catecholamine through a narrow fusion pore (van Kempen et al., 2011; Zhou et al., 1996). However, most slow FFN events last longer (up to 10-fold) than stand-alone feet, indicating the extreme stability of the narrow fusion pore. Monitoring chromaffin granules before exocytosis, we find that slow events derive exclusively from docked vesicles. In contrast, FFN release is invariably fast for events without a detectable docked vesicle. This may reflect a requirement of slow events for stable docking, or simply an inability to detect slow release from vesicles slightly above the TIRF plane.

### Dual Imaging of FFN and Peptide Release

The pH-insensitivity of FFN206 raises the possibility that slow FFN events may arise from vesicle movement within the TIRF plane rather than exocytosis. To monitor opening of the fusion pore by an independent method, we labeled chromaffin granules by expressing brain-derived neurotrophic factor (BDNF) fused to the ecliptic pHluorin, a highly pH-sensitive form of GFP (Miesenböck et al., 1998). BDNF-pHluorin sorts to DCVs where it is quenched by the low lumenal pH. Opening of the fusion pore exposes lumenal pHluorin to the neutral external pH, increasing its fluorescence. FFN and pHluorin are excited at different wavelengths, enabling us to image both fluorophores at the same events (Figure 2C–G and Figure 1—video 5). The weakly fluorescent BDNF-pHluorin required longer exposure times, reducing the rate of image acquisition and resulting in the loss or truncation of FFN fluorescence increase associated with fast events (Figure 2D,F,G). By dual imaging, we find that the vast majority of FFN events are accompanied by exocytosis of BDNF-pHluorin (147/166, Figure 2C). BDNF-pHluorin unquenching invariably follows release of the much smaller FFN206 (Figure 2D–G), due at least in part to the effects of lumenal buffering (Holz, 1978). A small number of FFN events are not associated with BDNF-pHluorin unquenching (19/166, Figure 2C), presumably reflecting FFN loaded into vesicles made before exogenous expression of BDNF-pHluorin. A fraction of BDNF-pHluorin events is not associated with FFN release (66/213, Figure 2C), indicating that FFN206 does not label all chromaffin granules.

Fast and slow events are accompanied by BDNF-pHluorin exocytosis, confirming that they are both exocytic, consistent with their dependence on stimulation, external Ca^++^, and v-SNARE. Corelease of BDNF-pHluorin also shows that FFNs release predominantly if not exclusively from peptidergic DCVs. Thus, the full range of FFN events derive from DCVs, and we do not need to invoke another class of vesicles to account for this variation.

### The Fusion Pore Can Limit Entry of External Dye

Since lumenal buffering delays the unquenching of pHluorin reporters (Holz, 1978), the precise moment when the fusion pore opens in slow FFN events remains unclear. We therefore monitored the entry of external Alexa fluor 488, a small, cell-impermeant fluorophore, as an alternative reporter for fusion pore opening (Chiang et al., 2014; Zhao et al., 2016). Chromaffin cells were loaded with FFN206 and bathed in solution containing Alexa Fluor 488 to monitor both FFN release and Alexa dye entry at the same events by simultaneous, dual imaging. The vast majority of FFN events are accompanied by Alexa dye entry (144/179, Figure 3A–G), consistent with exocytosis. Events identified by Alexa dye entry alone, without FFN release, may represent endocytosis (6/150, Figure 3G), whereas FFN-only events (35/179, Figure 3G) may represent rapid vesicle collapse that prevents dye entry (Chiang et al., 2014). All fast FFN events coincide with simultaneous Alexa dye entry (Figure 3A), as expected. For slow events detectable with both reporters, however, we were surprised to find a difference in timing between the onset of FFN fluorescence increase and Alexa dye entry (Figure 3B–F). In some cases, the FFN event and Alexa entry occur over the same extended period (Figure 3B). More often, dye entry lags the onset of FFN signal increase (Figures 3C,D) and in some cases occurs only toward the end of FFN discharge (Figure 3E,F).

**Figure 3.**
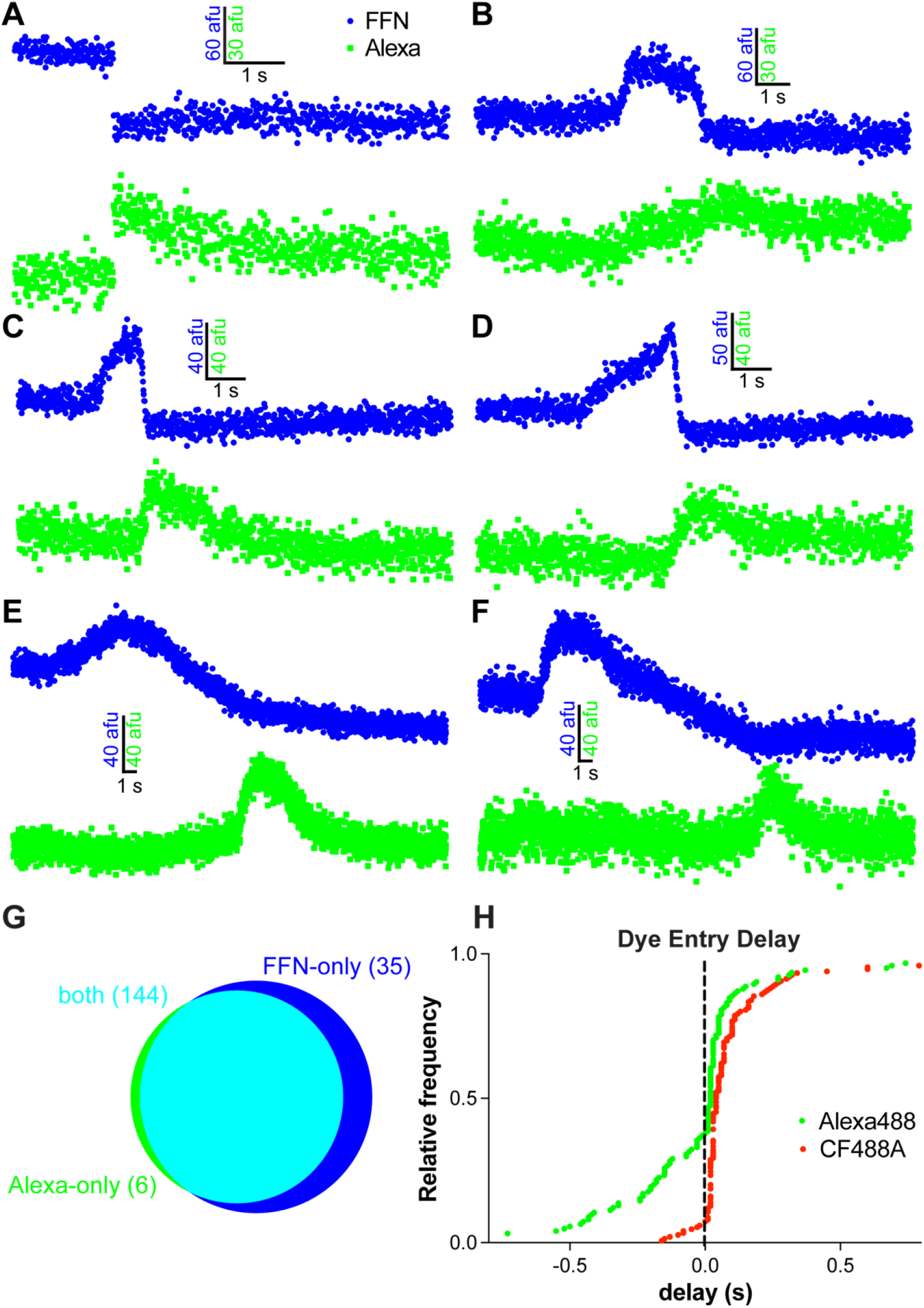
FFN and Alexa fluor 488 dual imaging. (**A–F**) Fluorescence time traces showing Alexa (green) dye entry occurs simultaneously with a fast FFN event (blue) (**A**) but during (**B**) or after a slow FFN event (**C–F**). (**G**) Quantitative Venn diagram showing that the vast majority of FFN events are accompanied by Alexa dye entry. (**H**) Onset of external dye entry relative to peak FFN fluorescence. Negative delay means external dye entry occurred before FFN reached peak fluorescence. **Figure 3—figure supplement 1.** Onset of external Alexa 488 and ATTO 655 entry relative to peak FFN206 fluorescence. **Figure 3—figure supplement 2.** Slow FFN release and Alexa dye entry imaged by confocal microscopy.

Figure 3H shows the temporal relationship between the onset of Alexa dye entry and peak FFN fluorescence. Although Alexa dye entry often follows the peak of FFN fluorescence, it can precede the peak by over 500 ms. The fusion pore can thus open during the gradual increase of FFN fluorescence, well before the peak. Since slow FFN events are exocytic, the decrease of FFN fluorescence after the peak must reflect release, requiring that the fusion pore open at or before the peak of FFN fluorescence.

What accounts for the delay in external Alexa dye entry? One possibility is restriction by the size of the fusion pore. Initially, the small size of the pore may exclude Alexa fluor 488 (MW 643 Da, Stokes radius 0.51 nm) but permit release of FFN206 (MW 219 Da, Stokes radius 0.30 nm), which is one third the MW and 60% the radius of the Alexa dye. Alexa dye entry may therefore require pore expansion, resulting in a delay relative to FFN release. This mechanism predicts that a dye larger than Alexa fluor 488 will enter with greater delay. Indeed, the larger CF488A (MW 914 Da, Stokes radius 0.61 nm) enters with even longer delay. The extent to which dye entry can precede peak FFN fluorescence is also reduced for CF488A (Figure 3H). These pore sizes are slightly smaller than the 1–2 nm reported previously (Llobet et al. 2008).

Alternatively, it has been reported that the fusion pore is cation-selective (Delacruz et al., 2021). Since both Alexa 488 and CF488A are anionic, it is possible that the delay in external dye entry may be due to electrostatic repulsion. However, we could not directly test cation selectivity due to the lack of commercially available, cationic dye that is cell impermeant. Instead, we compared the entry of Alexa 488 with ATTO 655, a zwitterionic dye that has no net charge at physiological pH and is similar to Alexa 488 in size (MW 634 Da). The entry of ATTO 655 is slightly more delayed than that of Alexa 488 (Figure 3—figure supplement 1), which is inconsistent with delay by electrostatic repulsion. Thus, it appears that the size of external dye is the main determinant of delay, with charge playing a secondary role.

### The Fluorescence Increase of Slow Events is Not Due to Movement

What accounts for the gradual increase of FFN fluorescence in slow events? Since the strength of the evanescent field increases exponentially with approach to the coverslip, the increase in fluorescence could reflect either vesicle movement or FFN release. In either of these cases, the fluorescence should not increase with excitation by a field of uniform strength. To test this idea, we imaged FFN events at the lateral edge of the cell by confocal microscopy. Using Alexa dye entry, we identified 25 events from nine cells as exocytic. Twenty of these events showed slow FFN release, and all of these exhibited an increase in fluorescence similar to that observed by TIRF microscopy (Figure 3—figure supplement 2). The increase in fluorescence for slow events thus cannot be explained by FFN movement, either within vesicles or by release. Vesicle swelling provides an alternative explanation: Since FFN206 self-quenches at high concentrations (Figure 1—figure supplement 1E), an increase in DCV volume would lower FFN concentration, relieve self-quenching and thereby increase fluorescence. The osmotic gradient across the chromaffin granule membrane (Sudhof, 1982) can provide the force that would drive this swelling, and both mast cell and chromaffin granules swell upon exocytosis (Breckenridge and Almers, 1987; Shin et al., 2020; Terakawa et al., 1991). Taken together, these observations suggest that vesicle fusion occurs at the onset of fluorescence increase for both slow and fast events.

### Quantitative Analysis of FFN Event Kinetics Reveals Distinct Modes of Fusion

High-speed imaging with FFN206 captures events with diverse fluorescence profiles. Some exhibit a characteristic spike whereas the rest vary widely, suggesting distinct mechanisms. To distinguish between this possibility and the prevailing assumption of a single, common mechanism with stochastic variation, we characterized the kinetics of all FFN events by their full width at half-maximum (FWHM), a simple metric broadly applicable to any curve with a peak that increases monotonically before and decreases monotonically after the maximum. However, variation in event profile required different methods to calculate FWHM. Events with a spike were fit by non-linear least-squares regression, whereas those without spikes were fit using free-knot B-splines (Dung and Tjahjowidodo, 2017). FWHM was then computed from the fitted curves (Figure 4A,B). To avoid classifying individual events into discrete categories, which prevents comparison across categories, we modeled the FWHM of individual events as a continuous random variable (Figure 4C). With no *a priori* classification, maximum likelihood estimation shows that the observed data are best explained by a mixture of two lognormal distributions (Figure 4D,E). The analysis of event half-life yields similar results (Figure 4—figure supplement 1). This is inconsistent with stochastic variation as the sole source of fusion pore heterogeneity and strongly suggests two distinct mechanisms for exocytosis.

**Figure 4.**
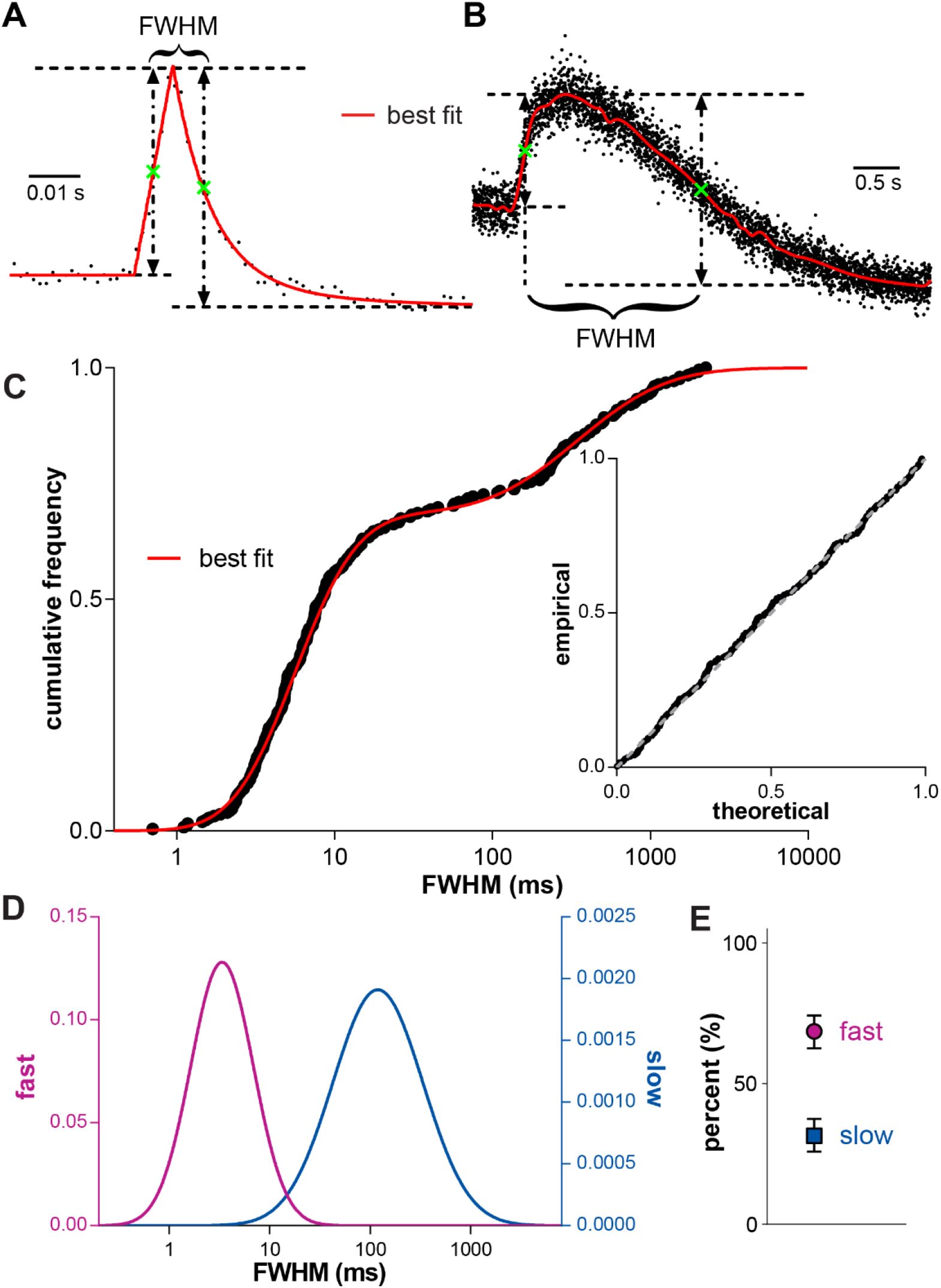
Quantitative analysis of FFN events. (**A**) Events with a spike were fit by least-squares regression. (**B**) Events without spikes were fit using free-knot B-splines. Event kinetics are characterized using the full width at half-maximum (FWHM), defined as the time difference between the half-rise and half-decay points (green crosses). (**C**) The observed cumulative frequency of FWHM for 252 events from 12 cells and the cumulative probability function determined by fitting the data using maximum likelihood estimation (MLE). Inset: P-P plot showing the goodness of fit. (**D**) The probability density functions of the two lognormal distributions used to fit the data in **C**. (**E**) Relative contribution of the two components shown in **D**. Error bars: 95% confidence interval. **Figure 4—figure supplement 1.** Characterizing FFN release kinetics using full width at half maximum.

Previous work has shown that membrane tension is a primary determinant of fusion pore behavior (Amatore et al., 2000; Bretou et al., 2014). We also observed that the enzyme used to digest extracellular matrix (ECM) of the adrenal gland when isolating chromaffin cells and the substrate used for cell attachment can have profound effects on exocytosis occurring within the surface of the cell attached to the substrate (TIRF footprint). Extensive removal of ECM with papain or the exclusive use of air-dried polylysine for substrate attachment can both result in tight adherence that eliminates FFN events within the TIRF footprint (data not shown). To determine whether local differences in substrate attachment account for bimodal fusion pore behavior, we again used the intensity of Alexa dye fluorescence in the TIRF field as a proxy for tightness of attachment. We found that FFN release kinetics do not correlate with the local tightness of attachment at the sites of fusion (correlation coefficient −0.075, Figure 1—figure supplement 3E). Thus, the tightness of substrate attachment does not account for bimodal behavior of the chromaffin granule fusion pore.

### Differential Regulation of Fast and Slow Events by Synaptotagmin 7

A number of proteins have been shown to influence the rate of fusion pore dilation (Anantharam et al., 2011; Graham et al., 2002; Doreian et al., 2008; Berberian et al., 2009; Han et al., 2004; Logan et al., 2017). These include the synaptotagmins, Ca^++^ sensors for regulated exocytosis (Sudhof, 2013). Synaptotagmin (syt) 1 and 7 are the major Ca^++^ sensors in chromaffin cells (Schonn et al., 2008), and loss of either isoform slows dilation of the chromaffin granule fusion pore, promoting kiss-and-run exocytosis (Das et al., 2020; Lynch et al., 2008; Segovia et al., 2010; Wang et al., 2001; Zhang et al., 2010). Syt 7 alone is responsible for ∼50% of all regulated exocytosis in chromaffin cells (Schonn et al., 2008), and accumulating evidence has suggested that it localizes to a distinct subset of chromaffin granules, conferring slow pore dilation (Bendahmane et al., 2018; Bendahmane et al., 2020; Jaiswal et al., 2004; Rao et al., 2014; Zhang et al., 2011).

To determine whether the two classes of fusion pore behavior depend on different Ca^++^ sensors, we compared FFN release kinetics in chromaffin cells from wild-type and syt 7 knockout (KO) mice. The cumulative frequency distribution of event FWHM differs significantly between wild-type and syt 7 KO cells (p < 10^−5^ by Kolmogorov-Smirnov) with an overall slowing of FFN release in the KO (Figure 5A), consistent with previous reports. Using maximum likelihood estimation, we then examined the two kinetic components of fusion pore behavior. Surprisingly, wild-type and KO show no significant difference in the kinetics of the corresponding components (Figure 5B). However, the proportion of each component differs significantly between wild-type and KO (Figure 5C). Wild-type cells exhibit ∼70% fast events and ∼30% slow, consistent with earlier observations (Figure 4E). In contrast, fast and slow events each account for ∼50% of the events in syt 7 KO cells, and the results do not change with analysis using half-life (Figure 5— figure supplement 1). We conclude that syt 7 alters the proportion of fast and slow events without changing their intrinsic properties. Thus, syt 7 plays a regulatory rather than direct role in fusion pore behavior. The two classes of fusion pore behavior respond differently to syt 7, supporting the involvement of distinct mechanisms and indicating the potential for independent regulation.

**Figure 5.**
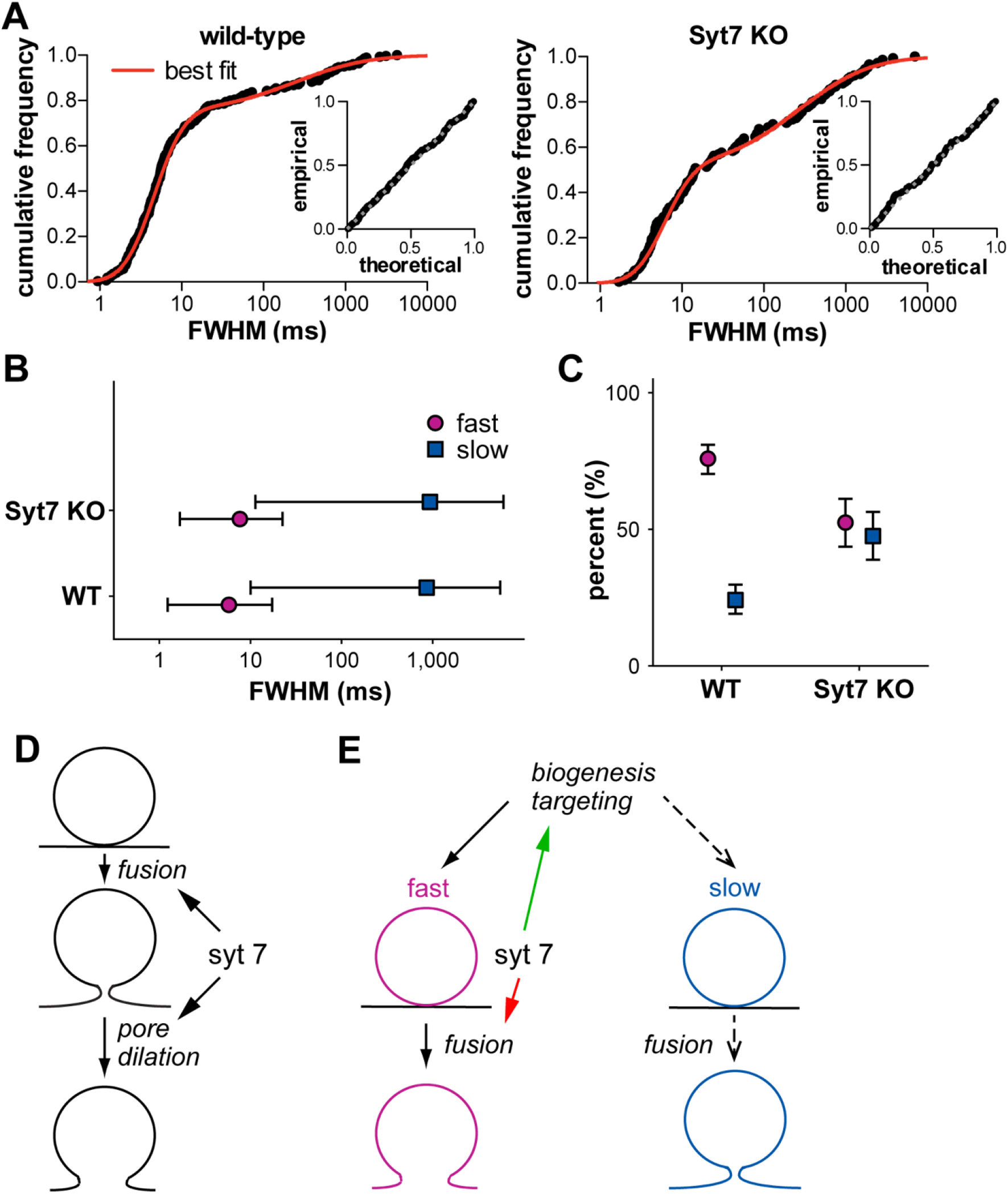
Synaptotagmin 7 knockout increases the proportion of slow events without affecting their kinetics. (**A**) Cumulative frequencies and MLE best fit of event full width half maximum (FWHM) in wild-type (left, 281 events from 14 cells) and syt 7 knockout (right, 144 events from 10 cells). Insets show the respective P-P plots. (**B**) Loss of syt 7 does not alter the properties of individual kinetic components. Error bars represent the 2.5% to 97.5% quantile region of each distribution, calculated using best-fit parameters. (**C**) Loss of syt 7 significantly reduces the relative contribution of the fast component to the total. Error bars indicate 95% confidence interval. (**D**) Traditional view that syt 7 mediates both membrane fusion and pore expansion. (**E**) The results suggest two distinct pathways to exocytosis, leading to fast (solid black arrows) or slow release (dashed arrows). Syt 7 may either act to promote the targeting of vesicles for fast release (green arrow), or to increase specifically the fusion probability of vesicles forming large pores (red arrow). **Figure 5—figure supplement 1.** The effects of synaptotagmin 7 knockout measured using full width at half maximum.

## DISCUSSION

The use of FFNs has enabled us to image DCV exocytosis at millisecond resolution. Catalytic accumulation of FFN within chromaffin granules increases the fluorescent signal, reducing the required exposure time. The small size of FFNs also makes their release kinetics a direct measure of fusion pore opening, unlike pHluorin unquenching which is delayed by dissolution of the dense core and lumenal buffering (Holz, 1978). In addition, the pH insensitivity of FFN206 enables detection of vesicle docking before fusion, in contrast to amperometry, which monitors release, and pHluorin imaging, in which the fluorophore is quenched before exocytosis. Taking advantage of these properties, we observe a range of FFN events from fast to slow, and all of these depend on stimulation, external Ca^++^ and v-SNARE. Moreover, all event types associate with the unquenching of BDNF-pHluorin and external dye entry, consistent with an exocytic mechanism.

The majority of FFN events derive from docked vesicles. A small number of vesicles appear to fuse immediately upon arrival at the plasma membrane. However, the apparently instantaneous docking, priming and fusion are inconsistent with previous observations (Chen et al., 2013; Steyer et al., 1997), and we cannot exclude fusion just outside the TIRF plane. We also observe very few events without a previously detected vesicle, consistent with a large population of stably docked granules.

All the observed events are exocytic, but when does the fusion pore open? For fast events, the rapid increase in fluorescence reflects bulk release of FFNs into the extracellular space, indicating that the fusion pore must open at the onset of fluorescence increase. For slow events, we observe a gradual increase in FFN fluorescence by confocal as well as TIRF microscopy, excluding vesicle or dye movement in the evanescent field. What then accounts for the increase in fluorescence of slow events? DCVs of both mast cells and chromaffin cells have been shown to swell after membrane fusion (Breckenridge and Almers, 1987; Terakawa et al., 1991), and FFN206 self-quenches at high concentrations, suggesting that vesicle swelling relieves FFN self-quenching, and that membrane fusion precedes the increase in FFN fluorescence. Recent work has indeed identified a form of chromaffin granule fusion involving slow release with vesicle enlargement (Shin et al., 2020) that presumably corresponds to the slow events reported here.

We also observe that for slow events, external dye enters considerably later than the onset of fluorescence increase. What causes this delay? We show that the extent of delay depends on the molecular weight of the external dye, suggesting occlusion by a narrow fusion pore. For slow events, the narrow pore also slows the exit of FFN. After opening, the fusion pore likely expands to accelerate FFN efflux and allow external dye entry. Larger external dyes enter later, as the pore expands over time. Previously, a delay in external dye entry was interpreted as evidence for a stable, hemifused state preceding full fusion (Chiang et al., 2014; Zhao et al., 2016). Our results suggest that this may instead reflect an extremely narrow pore initially less than 1 nm in diameter.

The ability to monitor the duration of all exocytic events makes the full distribution of fusion pore behavior amenable to quantitative methods. Previous work assigning exocytic events to different classes based on the presence or absence of certain features (Chiang et al., 2014; Logan et al., 2017; van Kempen et al., 2011) made it difficult to determine the relationship between classes and to assess the mechanistic basis for differences between them. In this work, we use a simple kinetic parameter applicable to all events, FWHM. By treating event FWHM as a continuous random variable, the analysis accounts for the inherent stochasticity of biological systems while avoiding the *a priori* classification of individual events. The analysis of FWHM distribution reveals two lognormal components, indicating two distinct modes of release that are inconsistent with stochastic variation in a common pathway.

What accounts for the difference between the two classes of release event? A possibility is that one of them does not involve a chromaffin granule. However, exocytosis of a peptide reporter accompanies both event classes, indicating that both involve DCVs. The local tightness of attachment also appears to have no role in the variability of event kinetics. Loss of syt 7 reduces the average rate of release, in agreement with some previous studies (Li et al., 2009; Segovia et al., 2010; Tsuboi and Fukuda, 2007; Zhang et al., 2010) but not all (Bendahmane et al., 2018; Bendahmane et al., 2020; Rao et al., 2014; Zhang et al., 2010; Zhang et al., 2011). This has generally been interpreted as evidence of a direct role for syt 7 in pore dilation as well as membrane fusion (Figure 5D). However, we find that the loss of syt 7 slows overall release by increasing the proportion of DCVs that undergo exocytosis with small pores (slow events), suggesting a role in the regulation of exocytosis rather than direct participation at the fusion pore. Differences in fusion pore behavior may reflect the exocytosis of different DCVs. Recent work has indeed suggested the segregation of syt 1 and 7 to chromaffin granules of different size (Rao et al., 2014; Zhang et al., 2011). Syt 7 may thus play a role upstream of exocytosis, in DCV biogenesis or targeting to the release site, producing vesicles that undergo fast release (Figure 5E, green arrow) (Vevea et al., 2021). Alternatively, syt 7 may act during exocytosis to promote specifically the fusion of DCVs destined for fast release, a role consistent with its known function as a calcium sensor for exocytosis (Figure 5E, red arrow). It will now be important to identify the factors that confer the two modes of release and to understand their differential regulation by synaptotagmin and other proteins. The analysis of biological variation presented here thus provides a new perspective on the function of syt 7 and should also serve to guide future work on the molecular mechanisms that underlie the bimodal nature of regulated DCV exocytosis.

The demonstration of two distinct exocytic mechanisms suggests a biological role for the diversity of DCV fusion pore behavior which has important implications for the release of peptides, monoamines and other neuromodulators. A global change due to regulation of a common exocytic pathway would in principle affect all release events, making it difficult to regulate signaling independently over different time scales. In contrast, two modes of release enable differential control of temporally distinct signals. The fast events associated with rapid pore dilation release more cargo acutely. A narrow pore slows transmitter efflux and also enables the trapping of residual peptide cargo for subsequent release. Bimodal fusion pore kinetics thus confer the potential for independent regulation of fast versus slow release, adding a new dimension to the temporal control of signaling. The defect in fusion pore dilation that underlies impaired insulin release in type 2 diabetes (Collins et al., 2016) indicates the potential for subversion of this mechanism in disease. Indeed, mouse syt 7 KO and human syt 7 knockdown islet cells both show impaired insulin secretion (Dolai et al., 2016; Gustavsson et al., 2008), consistent with our observation that loss of syt 7 increases the proportion of slow events: a pore narrow enough to limit the release of FFN will prevent the release of peptide.

## ACKNOWLEDGMENTS

This work was supported by grants from the NIH (R01 NS109295) and the Michael J. Fox Foundation to R.H.E. We thank Todd Logan for developing the use of FFNs in chromaffin cells and members of the Edwards lab for helpful discussion and suggestions. We thank Wade Regehr for sending the synaptotagmin 7 knockout mice. We thank DeLaine Larson, Kari Herrington and SoYeon Kim at the UCSF Center for Advanced Light Microscopy for technical assistance and advice. The algorithm for Dual-View calibration was generously provided by Damien Jullié.

## AUTHOR CONTRIBUTIONS

Conceptualization: R.H.E.; Methodology: P.Z., R.H.E.; Software: P.Z.; Validation: P.Z.; Formal Analysis: P.Z., D.R.; Investigation: P.Z.; Resources: R.H.E.; Data Curation: P.Z.; Writing – Original Draft: R.H.E.; Writing – Review & Editing: P.Z., D.R. and R.H.E.; Visualization: P.Z. and R.H.E.; Supervision: R.H.E.; Funding Acquisition: R.H.E.

## METHODS

### Primary chromaffin cell culture

Primary mouse adrenal chromaffin cells were isolated as previously described (Logan et al., 2017), with slight modification. Briefly, adrenal glands were harvested from animals 4–6 weeks old and placed in ice cold Ca^++^-, Mg^++^-free (CMF) Hank’s balanced salt solution (HBSS, UCSF cell culture facility [CCF] media production) supplemented with 10 U ml^−1^ penicillin G and 10 μg ml^−1^ streptomycin (pen-strep, UCSF CCF media production). Adrenal medullae were isolated and digested in CMF-HBSS containing collagenase type I (2.9 mg ml^−1^, Worthington), bovine serum albumin (BSA) (3 mg ml^−1^, Sigma), DNase I (0.15 mg ml^−1^, Sigma), penicillin G (10 U ml^−1^), and streptomycin (10 µg ml^−1^) at 37° C for 40 min with shaking at 800 rpm in a thermomixer (Eppendorf). Digested medullae were resuspended and the enzymatic reaction quenched by addition of 10 volumes cold CMF-HBSS. Isolated chromaffin cells were collected by centrifugation at 300 *g* for 10 min at 4° C and resuspended in pre-warmed DMEM / Ham’s F-12 1:1 mix (UCSF CCF media production) supplemented with 10% FBS (HyClone), penicillin G (4 U ml^−1^), and streptomycin (4 μg ml^−1^). Cell suspensions were plated drop-wise onto chambered coverglass (μ-Slide, Ibidi) coated with 100 μg ml^−1^ collagen (type I, Corning), 50 μg ml^−1^ fibronectin (Innovative Research), and 1 μg ml^−1^ BSA, and incubated at 37° C / 5% CO2 for 45 min before addition of fresh culture media or lentiviral supernatant for overnight incubation. Medium was replaced the next morning.

### Lentiviral production

Low passage Lenti-X 293T cells (Clontech) were seeded onto 6-well plates coated with 0.1 mg ml^−1^ poly-L-lysine and transfected using FuGENE HD transfection reagent (Promega) with a mixture of third-generation lentiviral backbone vector pJHUMCS that encodes BDNF-pHluorin, as well as accessory plasmids pVSV-G, pREV, and pRRE. Cells were switched into chromaffin cell culture medium the next day. The culture medium was collected 48 h after transfection and cell debris sedimented by centrifugation at 1,000 *g* for 10 min at RT. The resulting lentiviral supernatant was either used immediately or stored at −80° C.

### Fluorescent false neurotransmitter (FFN) loading

Chromaffin cells were loaded with FFNs immediately before imaging. The culture medium was replaced with DMEM high glucose (UCSF CCF media production) containing 10 μM FFN (Abcam) filtered through a 0.2 µ filter (FFN loading buffer). Cells were incubated in FFN loading buffer at 37° C / 5% CO2 for 1 h before switching into modified Tyrode’s solution (in mM, 140 NaCl, 10 HEPES-NaOH, pH 7.4, 10 glucose, 4.5 KCl, 5 CaCl2, 1 MgCl2) for imaging. Exocytosis was evoked by adding an equal volume of stimulation buffer (in mM, 54.5 NaCl, 10 HEPES-NaOH, pH 7.4, 10 glucose, 90 KCl, 5 CaCl2, 1 MgCl2). For experiments using Alexa dyes, 60 µM Alexa Fluor 488 carboxylic acid, tris(triethylammonium) salt (Thermo Fisher) was included in modified Tyrode’s solution and stimulation buffer.

### Microscopy

Chromaffin cells were imaged at room temperature using an inverted microscope (Ti-E, Nikon). TIRF microscopy was done using a MLC400 monolithic laser combiner (Agilent Technologies), ZT405/488/561/640rpc dichroic filter (Chroma), and Plan Apo Lambda 100× 1.45 N.A. oil objective (Nikon). Rapid FFN imaging was done using ET455/50m emission filter (Chroma) and Orca Flash 4.0 C11440-22C sCMOS camera (Hamamatsu). Dual imaging of FFN and BDNF-pHluorin was done using ZET488/561/635m emission filter (Chroma) and iXon DU-897 EMCCD camera (ANDOR). Dual imaging of FFN with Alexa Fluor 488 (ThermoFisher) or CF488A (Biotium) was done using Dual-View filter cube (Optical Insights) mounted with T510lpxrxt beam splitter (Chroma), ET460/36m and ET545/40m emission filters (Chroma), and ORCA Flash 4.0 C11440-22C sCMOS camera (Hamamatsu). Confocal imaging of FFN and Alexa Fluor 488 was done using an integrated laser engine (ANDOR), Borealis CSU-W1 spinning disk (ANDOR), ZT405/488/561/640rpcv2 dichroic filter (Chroma), Plan Apo VC 100× 1.4 N.A. oil objective (Nikon), ZET405/488/561/635 emission filter (Chroma), and iXon DU-888 EMCCD camera (ANDOR). Cells were imaged in modified Tyrode’s solution and exocytosis was stimulated with 45 mM K^+^ by adding an equal volume of stimulation buffer (in mM, 54.5 NaCl, 10 HEPES-NaOH, pH 7.4, 10 glucose, 90 KCl, 5 CaCl2, 1 MgCl2).

### Dual-View calibration

Images acquired using the Dual-View (Optical Insights) beam-splitter were calibrated to minimize distortion between the two channels. The optical configurations of the beam-splitter were mechanically calibrated using TetraSpeck microspheres (0.1 µm, ThermoFisher) that fluoresce in both channels. To correct for the remaining small shifts, images were aligned using a modified version of a previously described algorithm (Taylor et al., 2011). Briefly, before each imaging session, a calibration image of TetraSpeck microspheres was captured. At least eight beads that appear in both channels were manually selected and the coordinates of their respective centers refined by 2-D Gaussian fit. A third-degree polynomial spatial transform was then used to align the corresponding centers of each bead in the two channels and the coefficients were calculated by interpolation. The calculated coefficients were then used for a third-degree polynomial spatial transform of the Alexa channel for the images collected in the subsequent imaging session.

### Data analysis

Individual exocytotic events were identified manually in ImageJ by placing 5×5-pixel regions of interest (ROIs) over the center of events. Average intensity profiles were extracted using the Time Series Analyzer plugin (V3) and further processed using Python. Peak detection was done using continuous wavelet transform (Nenadic and Burdick, 2005) and verified by manual inspection. Pre- and post-event baselines were estimated by analyzing consecutive 20-frame sliding windows for statistical equivalency, and refined by manual inspection. Events for which both pre- and post-event baselines were identified underwent further processing. The null hypothesis that the fluorescence of the pre- and post-event baselines are equivalent was tested using the two-tailed t-test, treating each imaging frame as an independent observation. If the null hypothesis was rejected, the vesicle was categorized as docked, otherwise as without detectable prior docking. If a peak was detected, the trace was fitted to six models (linear / exponential rise and linear / exponential / two-phase exponential decay) by non-linear least-squares minimization using the lmfit package (Newville et al., 2016), and the best-fit model chosen using the Akaike information criterion corrected for limited sample size (AICc). If no peak was detected, the trace was fitted using free-knot non-uniform B-splines, using a direct method to optimize knot placement (Dung and Tjahjowidodo, 2017). Goodness of fit was evaluated by testing the normality of the residuals and verified by manual inspection. For some events that were not well fitted by either approach, the trace was smoothed by taking the 11-frame walking average and the smoothed curve was interpolated using a cubic spline. Interpolation results were also verified by manual inspection. From the processed curves, the peak time point as well as the time points at which the trace reaches the mid-point of fluorescence increase (half-rise) and the mid-point of fluorescence decay (half-decay) were determined. The half-life is defined as the time from peak to half-decay, whereas the full-width at half-maximum (FWHM) was defined as the time from half-rise to half-decay. The cumulative frequency distribution of the half-life or FWHM was then obtained and fitted to a log-normal or two-component log-normal mixture distribution by maximum likelihood estimation. The best-fit model was selected by performing a likelihood-ratio test. The confidence intervals of the best-fit parameters were determined by finding the 95% likelihood ratio confidence interval. The mean and 95% confidence interval of each kinetic component were computed from the best-fit parameters.

### Evaluating cell attachment

The TIRF footprint of individual chromaffin cells, visualized by Alexa Fluor 488 fluorescence, was subtracted from the image of an empty field of view. Relative attachment was then normalized to the maximum pixel value of the resulting image, with 0.0 representing the loosest attachment (beyond the evanescent field) and 1.0 representing the tightest attachment (directly against the cover glass).

## SUPPLEMENTAL MATERIAL

### SUPPLEMENTAL FIGURES

**Figure 1—figure supplement 1.**
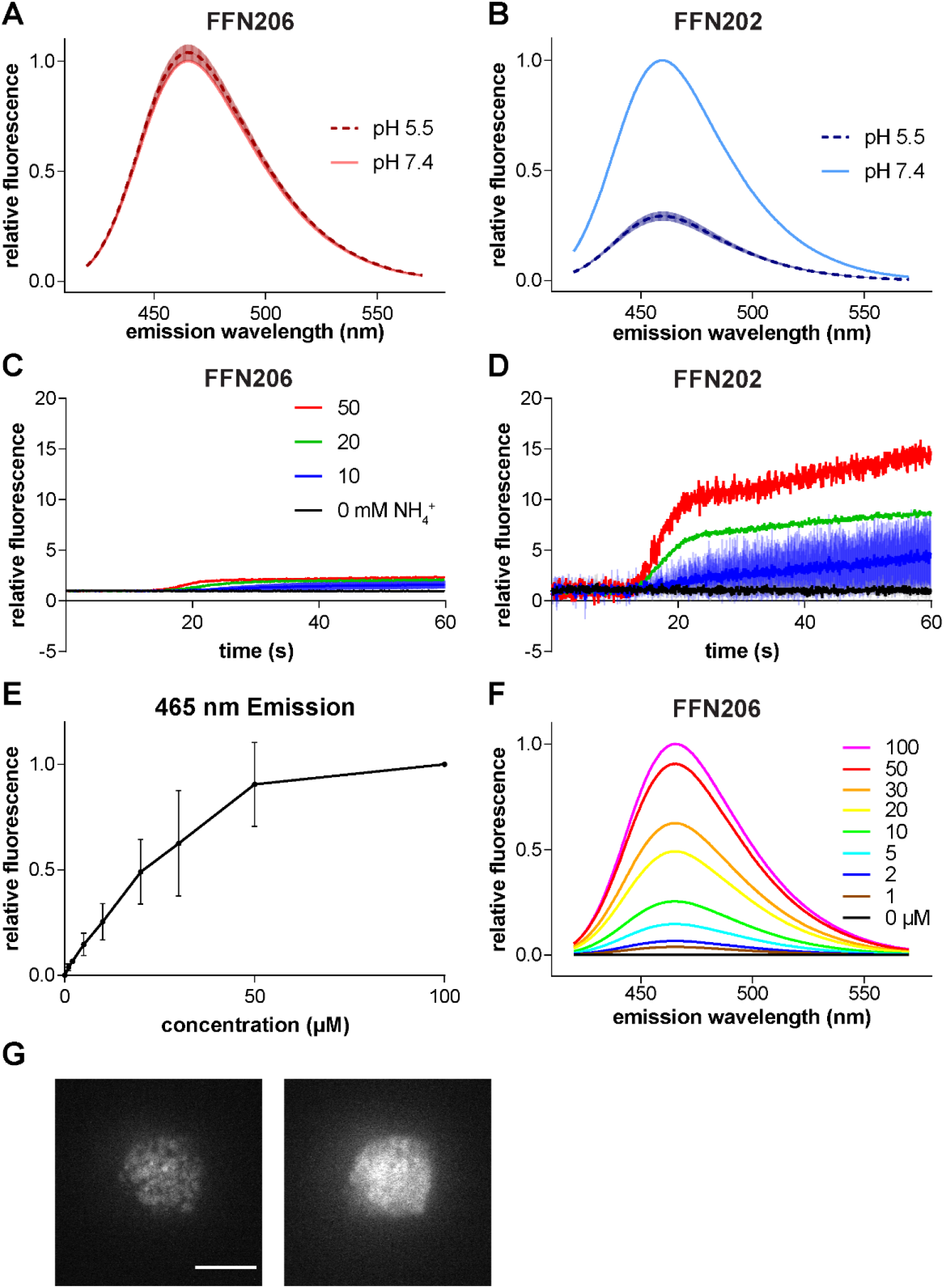
FFN properties. (**A–D**) pH sensitivity of FFN206 and FFN202. *In vitro* (**A** and **B**), FFN fluorescence was measured at 10 µM in pH 5.5 (dashed lines) and 7.4 (solid lines). Low pH does not affect FFN206 fluorescence (**A**), whereas FFN202 is quenched at pH 5.5 (**B**). In cells (**C** and **D**), FFN-loaded vesicles were unquenched by addition of NH4Cl. Cells loaded with FFN206 show much less unquenching (**C**) than cells loaded with FFN202 (**D**). (**E** and **F**) FFN206 exhibits self-quenching at higher concentrations. Measured *in vitro* at pH 7.4, FFN206 peak emission at 465 nm does not increase linearly with concentration (**E**). The emission spectrum does not change with concentration (**F**). (**G**) TIRF micrographs of an isolated primary chromaffin cell loaded with FFN206. Distinct FFN puncta are visible before addition of NH4Cl (left). Dissipation of the pH gradient with NH4Cl (50 mM) results in dispersion of the punctate fluorescence and an increase in overall fluorescence (right) presumably due to FFN206 leakage from vesicles and less self-quenching.

**Figure 1—figure supplement 2.**
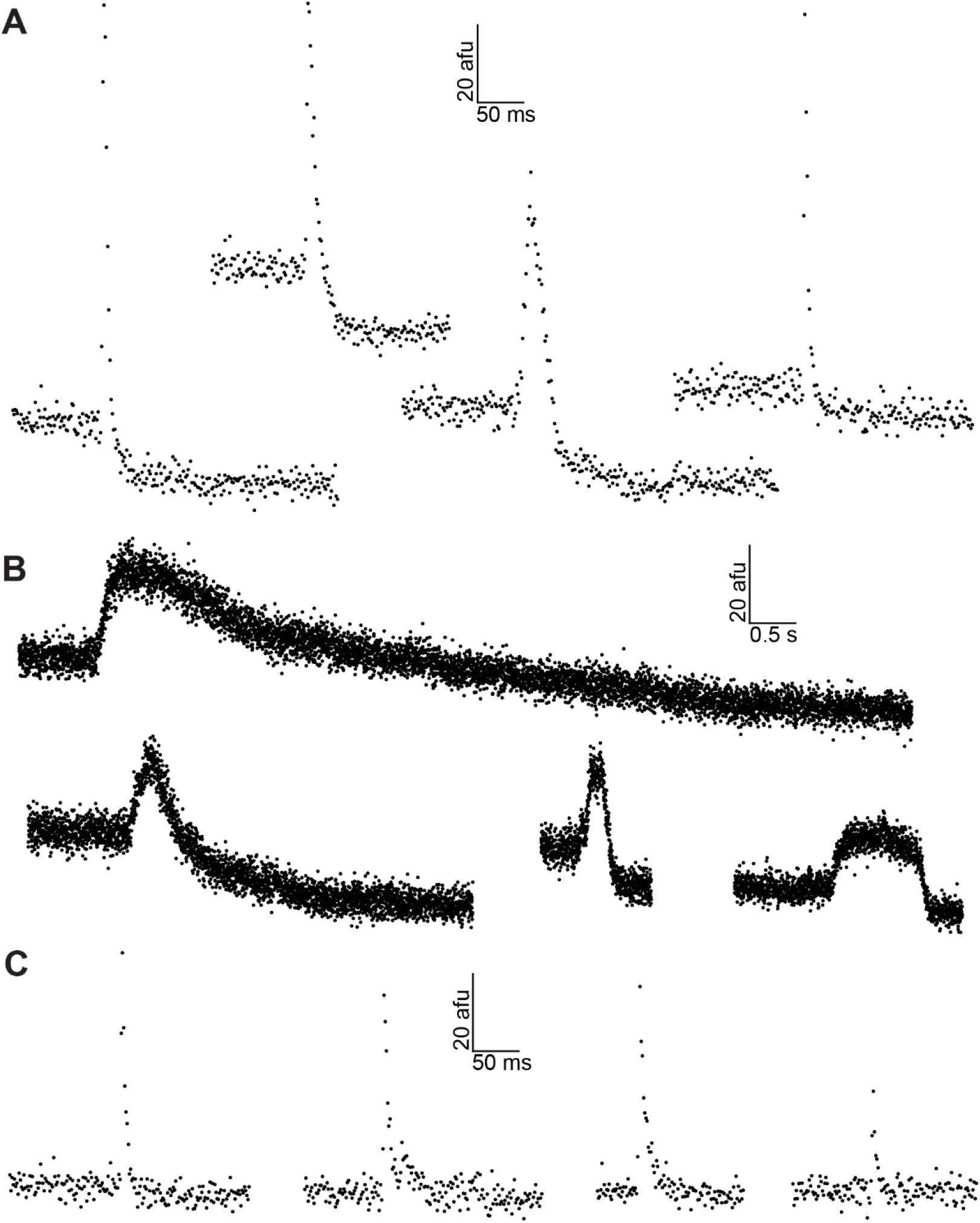
Sample fluorescence time traces of FFN release events. (**A–C**) FFN release events imaged by TIRF microscopy. Release events are broadly divided into three categories: rapid release with reduced post-event baseline (**A**), slow release with reduced post-event baseline (**B**), and rapid release with unchanged post-event baseline (**C**). Note the variation in event duration and profile among slow events.

**Figure 1—figure supplement 3.**
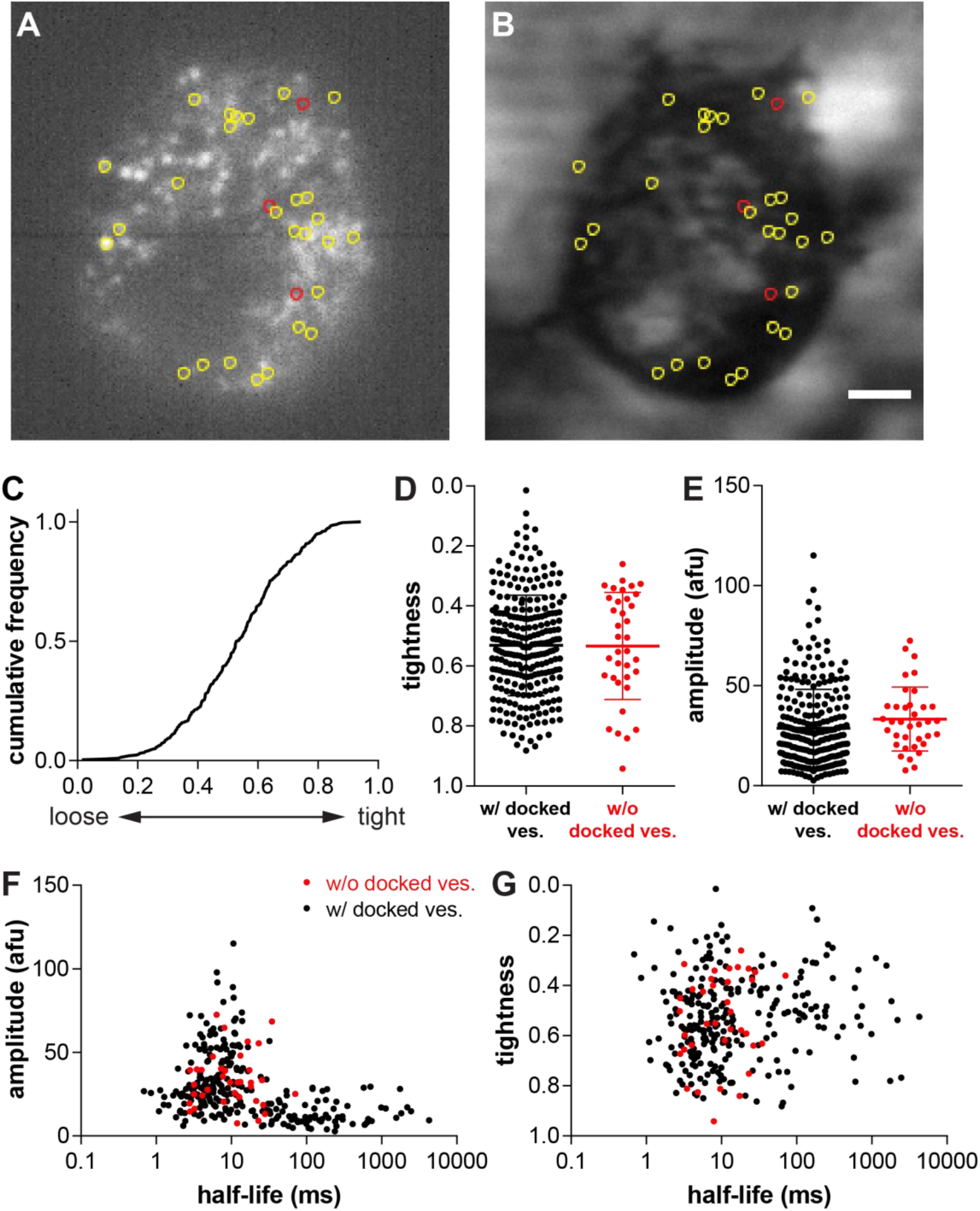
Tightness of cell attachment does not affect FFN release kinetics. (**A** and **B**) TIRF micrographs of an isolated primary mouse adrenal chromaffin cell overlaid with ROIs marking sites of FFN release, showing FFN (**A**) and Alexa Fluor 488 (**B**). The image in **A** shows the average of 10 frames, which is how events were identified (see Methods). Sites of FFN release with unchanged post-event baseline are indicated in red. The variation in background Alexa 488 fluorescence occurs over a larger spatial dimension likely reflecting unevenness of the glass coverslip. The brighter area to the upper right was observed in the same position relative to all cells and thus reflects refraction of the incident light. Scale bar: 2 µm. (**C**) Cumulative frequency distribution of relative attachment tightness at sites of FFN release (292 events from 21 cells). A relative tightness of 1.0 represents the tightest attachment (to the cover glass) whereas a value of 0.0 represents the loosest attachment (beyond the evanescent field). (**D**) Comparison of relative attachment tightness for sites with docked vesicles vs. sites without detectable docked vesicle. p = 0.7953 by two-tailed Mann-Whitney test. Error bars: standard deviation. (**E**) Comparison of peak amplitude for events with prior docked vesicles vs. those without detectable docked vesicle. p = 0.0286 by two-tailed Mann-Whitney test. Error bars: standard deviation. (**F**) Scatter plot showing the relationship between peak amplitude and FFN release kinetics. (**G**) Scatter plot showing the relationship between relative attachment tightness and FFN release kinetics. Covariance analysis yields a correlation coefficient of −0.008.

**Figure 3—figure supplement 1.**
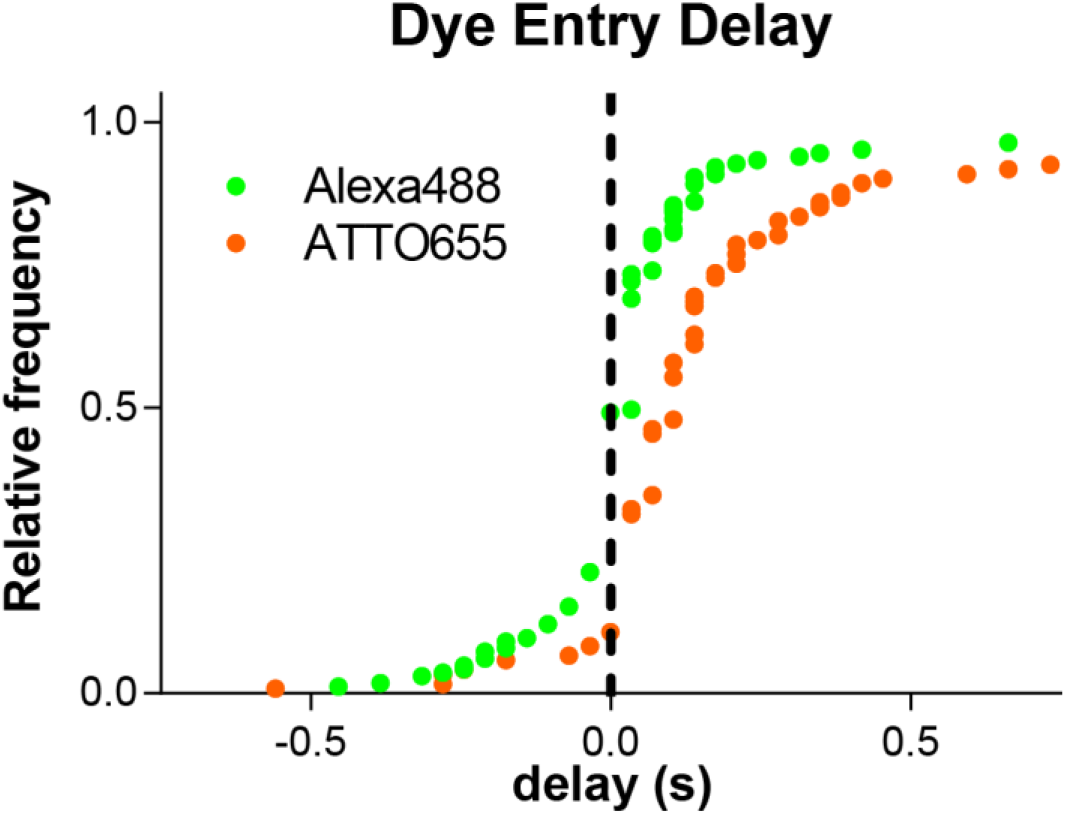
Onset of external Alexa 488 and ATTO 655 entry relative to peak FFN206 fluorescence. FFN206 release from chromaffin cells and entry of external dyes ATTO 655 or Alexa 488 were imaged in alternation, and the delay between FFN peak fluorescence and external dye entry measured. Negative delay means external dye entry occurred before the FFN reached peak fluorescence.

**Figure 3—figure supplement 2.**
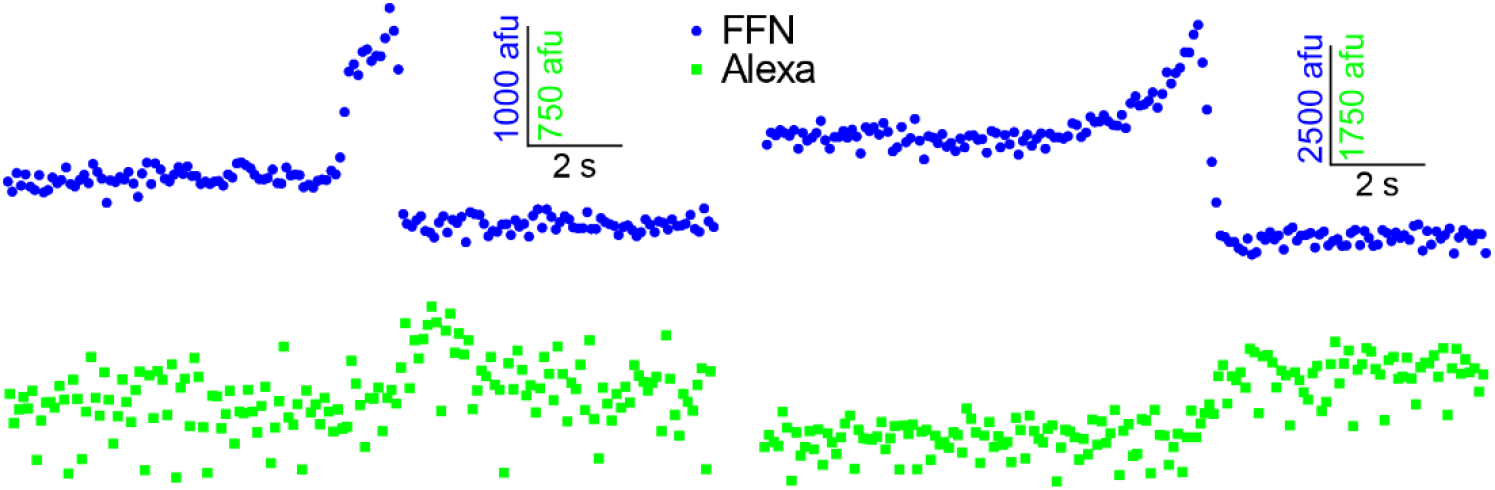
Slow FFN release and Alexa dye entry imaged by confocal microscopy. FFN206 release and Alexa dye entry were monitored in alternation by spinning-disk confocal microscopy. The confocal plane was set above the glass coverslip to image exocytic events that do not release toward the objective. FFN signal increases slowly before declining, similar to slow release events imaged by TIRF microscopy.

**Figure 4—figure supplement 1.**
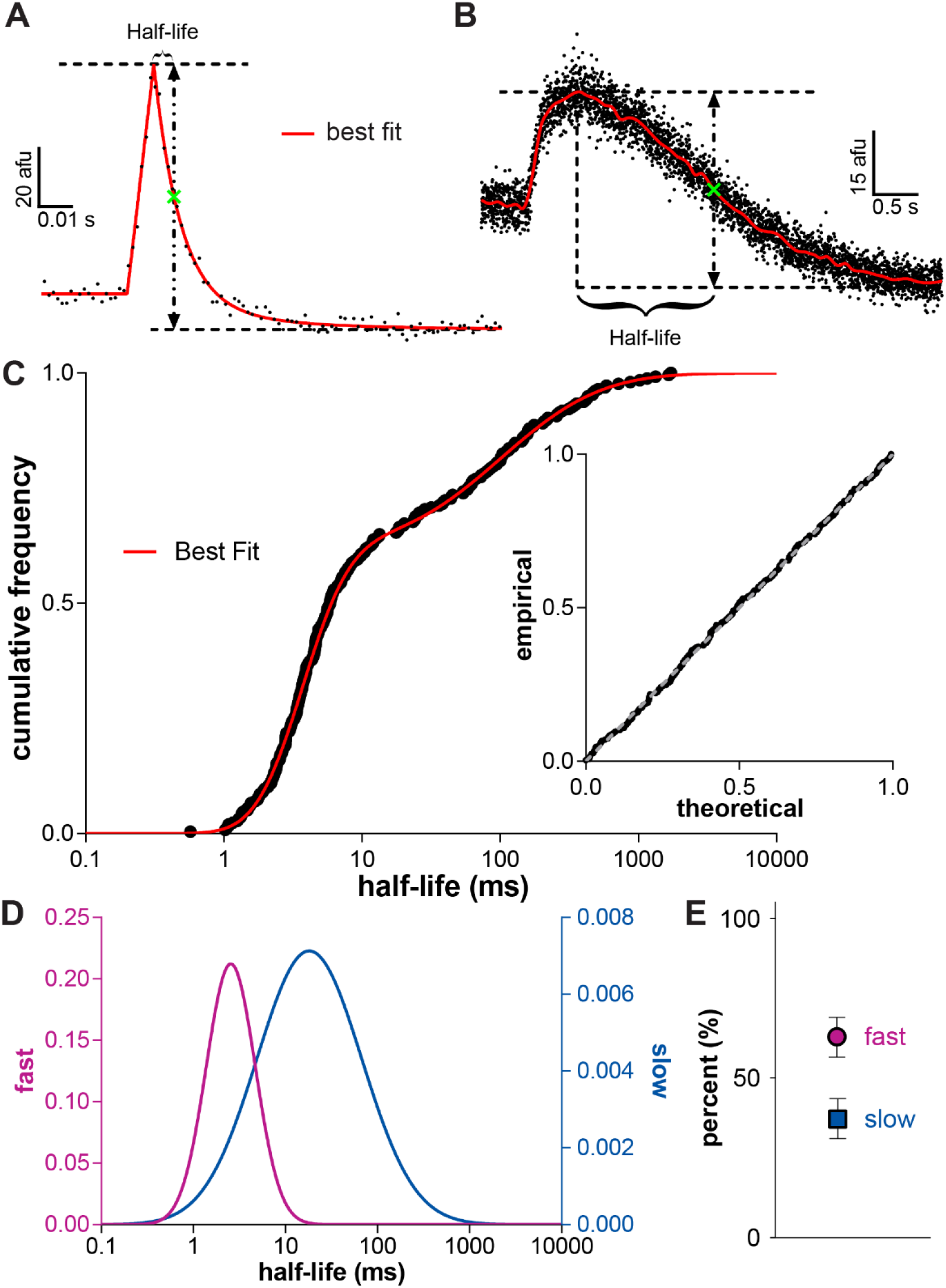
Characterizing FFN release kinetics using half-life. (**A** and **B**) The same FFN release data from Figure 3 was alternatively characterized using the half-life, defined as the time difference between peak and half-decay (green crosses). (**C**) The observed cumulative frequency of half-lives for 257 events from 12 cells in an experimental group, and the cumulative probability function determined by fitting the data using maximum likelihood estimation (MLE). Inset: P-P plot depicting the goodness of fit. (**D**) The probability density functions of the two lognormal distributions used to fit the data in (**C**). (**E**) Relative contribution of the two components shown in (D). Error bars, 95 % confidence interval.

**Figure 5—figure supplement 1.**
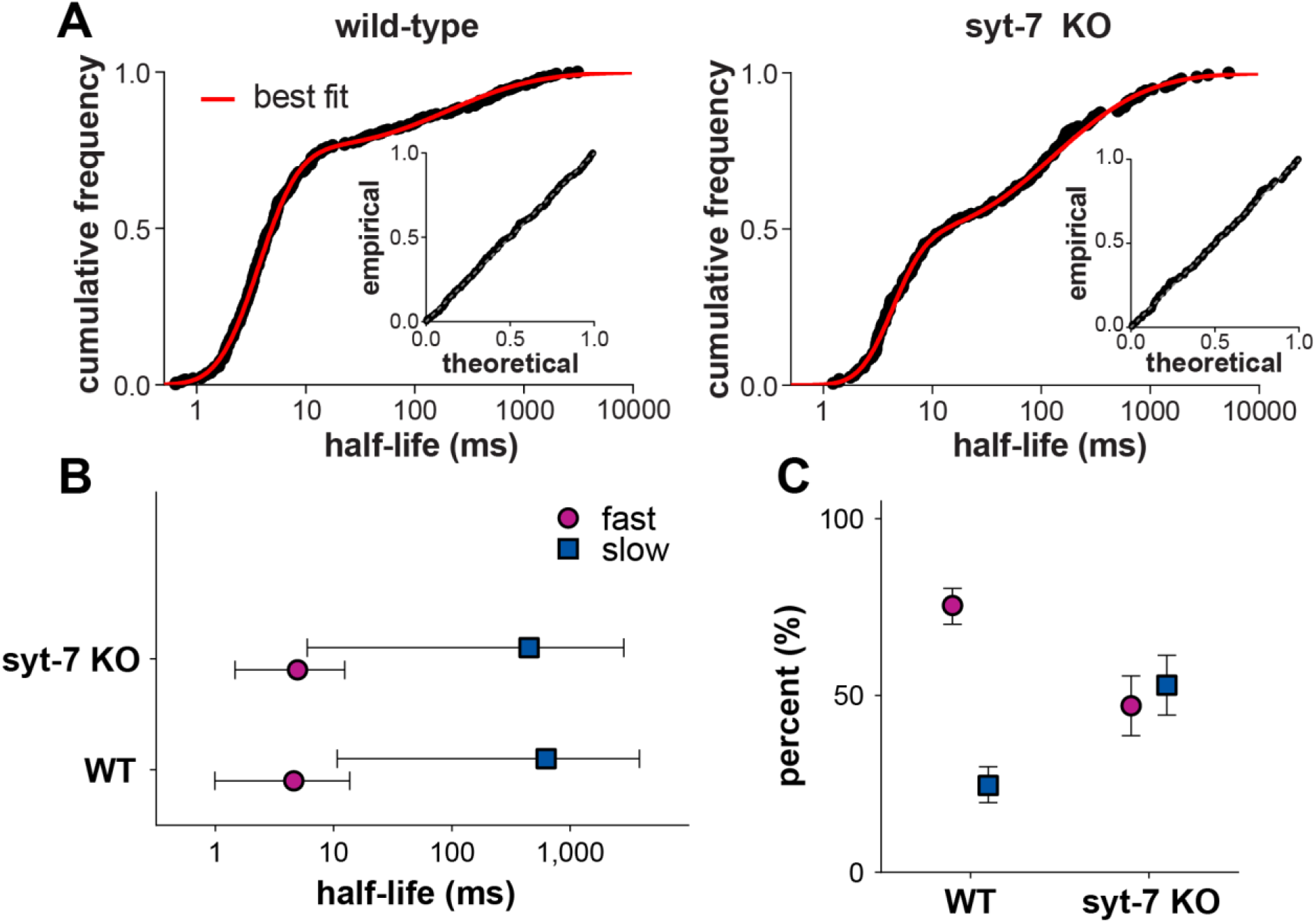
The effects of synaptotagmin 7 knockout measured using half-life. (**A**) Cumulative frequencies and MLE best fit of event half-lives in wild-type (left, 303 events from 14 cells) and syt 7 knockout (right, 157 events from 10 cells). Insets show the respective P-P plots depicting the goodness of fit. (**B**) Loss of syt 7 does not alter the properties of individual kinetic components. Error bars represent the 2.5 % to 97.5 % quantile region of each distribution, calculated using best-fit parameters. (**C**) Loss of syt 7 significantly reduces the relative contribution of the fast component to the total. Error bars indicate 95 % confidence interval.

**Figure 1—video 1. High-speed imaging of FFN release from an isolated primary mouse adrenal chromaffin cell in culture.**

After loading with FFN (10 µM) for one hour, the cell was imaged at rest for ∼5 s before stimulation with 45 mM K^+^. Imaging was performed at 800 Hz.

**Figure 1—video 2. Fast FFN release from a docked vesicle.**

An example event is marked by the arrow. The cell was loaded, stimulated and imaged as described for Video S1.

**Figure 1—video 3. Fast FFN release from an event without detectable prior docked vesicle.**

An example event is marked by the arrow. The cell was loaded, stimulated and imaged as described for Video S1.

**Figure 1—video 4. Slow FFN release from a docked vesicle.**

An example event is marked by the arrow. The cell was loaded, stimulated and imaged as described for Video S1.

**Figure 1—video 5. Dual imaging of FFN release and BDNF-pHluorin exocytosis.**

The cell was loaded as described for Video S1 and imaged at rest for ∼10 s before stimulation with 45 mM K^+^. The FFN channel is shown on the left and the BDNF-pHluorin channel on the right. The arrowhead marks a fast FFN release event in both channels. An arrow similarly marks a slow FFN event. (The slow event will be better visualized by increasing the frame rate.)

## REFERENCES

Alabi, A.A., and Tsien, R.W. (2013). Perspectives on kiss-and-run: role in exocytosis, endocytosis, and neurotransmission. Annu. Rev. Physiol. 75, 393–422.

Albillos, A., Dernick, G., Horstmann, H., Almers, W., Alvarez de Toledo, G., and Lindau, M. (1997). The exocytotic event in chromaffin cells revealed by patch amperometry. Nature 389, 509–512.

Amatore, C., Bouret, Y., Travis, E.R., and Wightman, R.M. (2000). Adrenaline release by chromaffin cells: constrained swelling of the vesicle matrix leads to full fusion at the ENS. Angew. Chem. Int. Ed. Engl. 39, 1952–1955.

Bao, H., Das, D., Courtney, N.A., Jiang, Y., Briguglio, J.S., Lou, X., Roston, D., Cui, Q., Chanda, B., and Chapman, E.R. (2018). Dynamics and number of trans-SNARE complexes determine nascent fusion pore properties. Nature 554, 260–263.

Bendahmane, M., Bohannon, K.P., Bradberry, M.M., Rao, T.C., Schmidtke, M.W., Abbineni, P.S., Chon, N.L., Tran, S., Lin, H., Chapman, E.R., et al. (2018). The synaptotagmin C2B domain calcium-binding loops modulate the rate of fusion pore expansion. Mol. Biol. Cell 29, 834–845.

Bendahmane, M., Morales, A., Kreutzberger, A.J.B., Schenk, N.A., Mohan, R., Bakshi, S., Philippe, J.M., Zhang, S., Kiessling, V., Tamm, L.K., et al. (2020). Synaptotagmin-7 enhances calcium-sensing of chromaffin cell granules and slows discharge of granule cargos. J. Neurochem. 154, 598–617.

Breckenridge, L.J., and Almers, W. (1987). Final steps in exocytosis observed in a cell with giant secretory granules. Proc. Natl. Acad. Sci. USA 84, 1945–1949.

Bretou, M., Jouannot, O., Fanget, I., Pierobon, P., Larochette, N., Gestraud, P., Guillon, M., Emiliani, V., Gasman, S., Desnos, C., et al. (2014). Cdc42 controls the dilation of the exocytotic fusion pore by regulating membrane tension. Mol. Biol. Cell 25, 3195–3209.

Chen, M., Van Hook, M.J., Zenisek, D., and Thoreson, W.B. (2013). Properties of ribbon and non-ribbon release from rod photoreceptors revealed by visualizing individual synaptic vesicles. J. Neurosci. 33, 2071–2086.

Chiang, H.C., Shin, W., Zhao, W.D., Hamid, E., Sheng, J., Baydyuk, M., Wen, P.J., Jin, A., Momboisse, F., and Wu, L.G. (2014). Post-fusion structural changes and their roles in exocytosis and endocytosis of dense-core vesicles. Nat. Commun. 5, 3356.

Chow, R.H., von Rüden, L., and Neher, E. (1992). Delay in vesicle fusion revealed by electrochemical monitoring of single secretory events in adrenal chromaffin cells. Nature 356, 60– 63.

Collins, S.C., Do, H.W., Hastoy, B., Hugill, A., Adam, J., Chibalina, M.V., Galvanovskis, J., Godazgar, M., Lee, S., Goldsworthy, M., et al. (2016). Increased Expression of the Diabetes Gene SOX4 Reduces Insulin Secretion by Impaired Fusion Pore Expansion. Diabetes 65, 1952–1961.

Das, D., Bao, H., Courtney, K.C., Wu, L., and Chapman, E.R. (2020). Resolving kinetic intermediates during the regulated assembly and disassembly of fusion pores. Nat. Commun. 11, 231.

Delacruz, J.B., Sharma, S., Rathore, S.S., Huang, M., Lenz, J.S., and Lindau, M. (2021). Fusion pores with low conductance are cation selective. Cell Rep. 36, 109580.

Dolai, S., Xie, L., Zhu, D., Liang, T., Qin, T., Xie, H., Kang, Y., Chapman, E.R., and Gaisano, H.Y. (2016). Synaptotagmin-7 Functions to Replenish Insulin Granules for Exocytosis in Human Islet beta-Cells. Diabetes 65, 1962–1976.

Dung, V.T., and Tjahjowidodo, T. (2017). A direct method to solve optimal knots of B-spline curves: An application for non-uniform B-spline curves fitting. PLoS One 12, e0173857.

Fulop, T., Radabaugh, S., and Smith, C. (2005). Activity-dependent differential transmitter release in mouse adrenal chromaffin cells. J. Neurosci. 25, 7324–7332.

Gubernator, N.G., Zhang, H., Staal, R.G., Mosharov, E.V., Pereira, D.B., Yue, M., Balsanek, V., Vadola, P.A., Mukherjee, B., Edwards, R.H., et al. (2009). Fluorescent false neurotransmitters visualize dopamine release from individual presynaptic terminals. Science 324, 1441–1444.

Gustavsson, N., Lao, Y., Maximov, A., Chuang, J.C., Kostromina, E., Repa, J.J., Li, C., Radda, G.K., Sudhof, T.C., and Han, W. (2008). Impaired insulin secretion and glucose intolerance in synaptotagmin-7 null mutant mice. Proc. Natl. Acad. Sci. USA 105, 3992–3997.

Holz, R.W. (1978). Evidence that catecholamine transport into chromaffin vesicles is coupled to vesicle membrane potential. Proc. Natl. Acad. Sci. USA 75, 5190–5194.

Hu, G., Henke, A., Karpowicz, R.J., Jr., Sonders, M.S., Farrimond, F., Edwards, R., Sulzer, D., and Sames, D. (2013). New fluorescent substrate enables quantitative and high-throughput examination of vesicular monoamine transporter 2 (VMAT2). ACS Chem. Biol. 8, 1947–1954.

Jaiswal, J.K., Chakrabarti, S., Andrews, N.W., and Simon, S.M. (2004). Synaptotagmin VII restricts fusion pore expansion during lysosomal exocytosis. PLoS Biol. 2, e233.

Kaeser, P.S., and Regehr, W.G. (2014). Molecular mechanisms for synchronous, asynchronous, and spontaneous neurotransmitter release. Ann. Rev. Physiol. 76, 333–363.

Li, J., Xiao, Y., Zhou, W., Wu, Z., Zhang, R., and Xu, T. (2009). Silence of Synaptotagmin VII inhibits release of dense core vesicles in PC12 cells. Sci. China C. Life Sci. 52, 1156–1163.

Lielig O, Baumgaertel R, and Bausch A.(2009) Selective filtering of particles by the extracellular matrix: an electrostatic band-pass. Biophys. J. 97, 1569–1577.

Llobet A, Wu M, and Langnado L (2008). The mouth of a dense-core vesicle opens and closes in a conceted action regulated by calcium and amphiphysin. J. Cell Biol. 182, 1017–1028,

Logan, T., Bendor, J., Toupin, C., Thorn, K., and Edwards, R.H. (2017). alpha-Synuclein promotes dilation of the exocytotic fusion pore. Nat. Neurosci. 20, 681–689.

Lynch, K.L., Gerona, R.R., Kielar, D.M., Martens, S., McMahon, H.T., and Martin, T.F. (2008). Synaptotagmin-1 utilizes membrane bending and SNARE binding to drive fusion pore expansion. Mol. Biol. Cell 19, 5093–5103.

McMahon, H.T., Ushkaryov, Y.A., Edelmann, L., Link, E., Binz, T., Niemann, H., Jahn, R., and Sudhof, T.C. (1993). Cellubrevin is a ubiquitous tetanus-toxin substrate homologous to a putative synaptic vesicle fusion protein. Nature 364, 346–349.

Miesenböck, G., De Angelis, D.A., and Rothman, J.E. (1998). Visualizing secretion and synaptic transmission with pH-sensitive green fluorescent proteins. Nature 394, 192–195.

Nenadic, Z., and Burdick, J.W. (2005). Spike detection using the continuous wavelet transform. IEEE Trans. Biomed. Eng. 52, 74–87.

Newville, M., Stensitzki, T., Allen, D.B., Rawlik, M., Ingargiola, A., and Nelson, A. (2016). LMFIT: non-linear least-square minimization and curve-fitting for Python. Astrophysics Source Code Library.

Parsons, T.D., Coorssen, J.R., Horstmann, H., and Almers, W. (1995). Docked granules, the exocytic burst, and the need for ATP hydrolysis in endocrine cells. Neuron 15, 1085–1096.

Perrais, D., Kleppe, I.C., Taraska, J.W., and Almers, W. (2004). Recapture after exocytosis causes differential retention of protein in granules of bovine chromaffin cells. J. Physiol. 560, 413– 428.

Rao, T.C., Passmore, D.R., Peleman, A.R., Das, M., Chapman, E.R., and Anantharam, A. (2014). Distinct fusion properties of synaptotagmin-1 and synaptotagmin-7 bearing dense core granules. Mol. Biol. Cell 25, 2416–2427.

Rorsman, P., and Renstrom, E. (2003). Insulin granule dynamics in pancreatic beta cells. Diabetologia 46, 1029–1045.

Schiavo, G., Benfenati, F., Poulain, B., Rossetto, O., Polverino de Laureto, P., DasGupta, B.R., and Montecucco, C. (1992). Tetanus and botulinum-B neurotoxins block neurotransmitter release by proteolytic cleavage of synaptobrevin. Nature 359, 832–835.

Schonn, J.S., Maximov, A., Lao, Y., Sudhof, T.C., and Sorensen, J.B. (2008). Synaptotagmin-1 and -7 are functionally overlapping Ca2+ sensors for exocytosis in adrenal chromaffin cells. Proc. Natl. Acad. Sci. USA 105, 3998–4003.

Segovia, M., Ales, E., Montes, M.A., Bonifas, I., Jemal, I., Lindau, M., Maximov, A., Sudhof, T.C., and Alvarez de Toledo, G. (2010). Push-and-pull regulation of the fusion pore by synaptotagmin-7. Proc. Natl. Acad. Sci. USA 107, 19032–19037.

Shin, W., Arpino, G., Thiyagarajan, S., Su, R., Ge, L., McDargh, Z., Guo, X., Wei, L., Shupliakov, O., Jin, A., et al. (2020). Vesicle Shrinking and Enlargement Play Opposing Roles in the Release of Exocytotic Contents. Cell Rep. 30, 421–431.e7.

Shin, W., Ge, L., Arpino, G., Villarreal, S.A., Hamid, E., Liu, H., Zhao, W.D., Wen, P.J., Chiang, H.C., and Wu, L.G. (2018). Visualization of membrane pore in live cells Reveals a dynamic-pore theory governing fusion and endocytosis. Cell 173, 934–945.e12.

Steyer, J.A., Horstmann, H., and Almers, W. (1997). Transport, docking and exocytosis of single secretory granules in live chromaffin cells. Nature 388, 474–478.

Stylianopoulos T, Poh M, Insin N, Bawendl MG, Fukumura D, Murin L, and Jain R. (2010) Diffusion of particles in the extracellular matrix: The effect of repulsive electrostatic interactions. Biophys. J. 99, 1342–1349.

Sudhof, T.C. (1982). Core structure, internal osmotic pressure and irreversible structural changes of chromaffin granules during osmometer behaviour. Biochim. Biophys. Acta 684, 27–39.

Sudhof, T.C. (2013). Neurotransmitter release: the last millisecond in the life of a synaptic vesicle. Neuron 80, 675–690.

Taylor, M.J., Perrais, D., and Merrifield, C.J. (2011). A high precision survey of the molecular dynamics of mammalian clathrin-mediated endocytosis. PLoS Biol. 9, e1000604.

Terakawa, S., Fan, J.H., Kumakura, K., and Ohara-Imaizumi, M. (1991). Quantitative analysis of exocytosis directly visualized in living chromaffin cells. Neurosci. Lett. 123, 82–86.

Tsuboi, T., and Fukuda, M. (2007). Synaptotagmin VII modulates the kinetics of dense-core vesicle exocytosis in PC12 cells. Genes Cells 12, 511–519.

van Kempen, G.T., vanderLeest, H.T., van den Berg, R.J., Eilers, P., and Westerink, R.H. (2011). Three distinct modes of exocytosis revealed by amperometry in neuroendocrine cells. Biophys. J. 100, 968–977.

Vevea, J.D., Kusick, G.F., Courtney, K.C., Chen, E., Watanabe, S., and Chapman, E.R. (2021). Synaptotagmin 7 is targeted to the axonal plasma membrane through gamma-secretase processing to promote synaptic vesicle docking in mouse hippocampal neurons. Elife 10.

Voets, T., Neher, E., and Moser, T. (1999). Mechanisms underlying phasic and sustained secretion in chromaffin cells from mouse adrenal slices. Neuron 23, 607–615.

Wang, C.T., Grishanin, R., Earles, C.A., Chang, P.Y., Martin, T.F., Chapman, E.R., and Jackson, M.B. (2001). Synaptotagmin modulation of fusion pore kinetics in regulated exocytosis of dense-core vesicles. Science 294, 1111–1115.

Xia, X., Lessmann, V., and Martin, T.F. (2009). Imaging of evoked dense-core-vesicle exocytosis in hippocampal neurons reveals long latencies and kiss-and-run fusion events. J. Cell Sci. 122, 75–82.

Zhang, Z., Hui, E., Chapman, E.R., and Jackson, M.B. (2010). Regulation of exocytosis and fusion pores by synaptotagmin-effector interactions. Mol. Biol. Cell 21, 2821–2831.

Zhang, Z., Wu, Y., Wang, Z., Dunning, F.M., Rehfuss, J., Ramanan, D., Chapman, E.R., and Jackson, M.B. (2011). Release mode of large and small dense-core vesicles specified by different synaptotagmin isoforms in PC12 cells. Mol. Biol. Cell 22, 2324–2336.

Zhao, W.D., Hamid, E., Shin, W., Wen, P.J., Krystofiak, E.S., Villarreal, S.A., Chiang, H.C., Kachar, B., and Wu, L.G. (2016). Hemi-fused structure mediates and controls fusion and fission in live cells. Nature 534, 548–552.

Zhou, Z., Misler, S., and Chow, R.H. (1996). Rapid fluctuations in transmitter release from single vesicles in bovine adrenal chromaffin cells. Biophys. J. 70, 1543–1552.

